# A dual-gene reporter-amplifier architecture for enhancing the sensitivity of molecular MRI by water exchange

**DOI:** 10.1101/2024.01.22.576672

**Authors:** Yimeng Huang, Xinyue Chen, Ziyue Zhu, Arnab Mukherjee

## Abstract

The development of genetic reporters for magnetic resonance imaging (MRI) is essential for investigating biological functions in intact animals. However, current MRI reporters have low sensitivity, making it challenging to create significant contrast against the tissue background, especially when only a small percentage of cells express the reporter. To overcome this limitation, we developed an approach that amplifies signals by co-expressing an MRI reporter gene, Oatp1b3, with a water channel, aquaporin-1 (Aqp1). We first show that the expression of Aqp1 amplifies the paramagnetic relaxation effect of Oatp1b3 by facilitating transmembrane water exchange. This mechanism provides Oatp1b3-expressing cells with access to a larger water pool compared with typical exchange-limited conditions. We further demonstrated that our methodology allows dual-labeled cells to be detected using approximately 10-fold lower concentrations of contrast agent than that in the Aqp1-free scenario. Finally, we show that our approach enables the imaging of mixed-cell populations containing a low fraction of Oatp1b3-labeled cells that are otherwise undetectable based on Oatp1b3 expression alone.

## Introduction

Investigation of in vivo biological functions and the development of gene- and cell-based therapies require the use of reporter genes to track cellular and molecular events in intact organisms. Reporters that rely on optical excitation have limited utility in this context because of the absorption and scattering of light by biological tissues^1,2^. Consequently, efforts have been made to create reporter genes compatible with tissue-penetrant techniques such as nuclear^3–6^, ultrasound^7–10^, and magnetic resonance imaging (MRI)^11–23^. Among these techniques, MRI offers the most detailed three-dimensional views of tissues, organs, and organ systems, with a high spatial resolution and no exposure to ionizing radiation. These unique advantages have motivated numerous efforts to develop genetically encoded MRI reporters^24,25^. Most biomolecular reporters for MRI rely on proteins that interact with paramagnetic metals, such as iron^11–14,21,26^, manganese^16,17,27^, and gadolinium^23,28–30^, to shorten the spin-lattice (T_1_) and spin-spin relaxation times (T_2_) of water molecules, allowing for the visualization of genetically labeled cells using T_1_- and T_2_-weighted imaging. In addition to paramagnetic relaxation, other mechanisms based on the diffusion of water molecules^20,22^ and chemical exchange of protons^15,31–35^ have also been explored. However, a major challenge for all classes of MRI reporter genes is their low sensitivity, which makes it difficult to create a significant contrast against the tissue background, especially when only a small population of cells expresses the reporter. This presents a significant obstacle in the application of MRI reporters to visualize biological processes such as neural activity, tumor metastasis, host-pathogen interactions, and cell trafficking, which often involve small numbers of reporter gene expressing cells.

To overcome the sensitivity limitations of current MRI reporters, we sought to develop an approach that combines paramagnetic relaxation enhancement with increased transmembrane water diffusion through the expression of two genetic constructs: an organic anion-transporting polypeptide, 1b3 (Oatp1b3), and a water channel, aquaporin-1 (Aqp1). Oatp1b3 is a hepatic drug transporter^36^ that functions as an MRI reporter, because it facilitates the uptake of gadolinium ethoxybenzyl diethylenetriamine pentaacetic acid (Gd-EOB-DTPA)^37^, a clinically approved contrast agent^23,28,38–40^. This results in shortening of the T_1_ of water molecules within Oatp1b3-expressing cells, thereby enabling their visualization through T_1_-weighted MRI (**Fig. 1a**). Aqp1, on the other hand, is a channel protein that facilitates the free and selective exchange of water molecules across the cell membrane and has been adapted as a reporter gene for visualizing cells using diffusion-weighted MRI^20,41^ (**Fig. 1b**).

**Figure 1:**
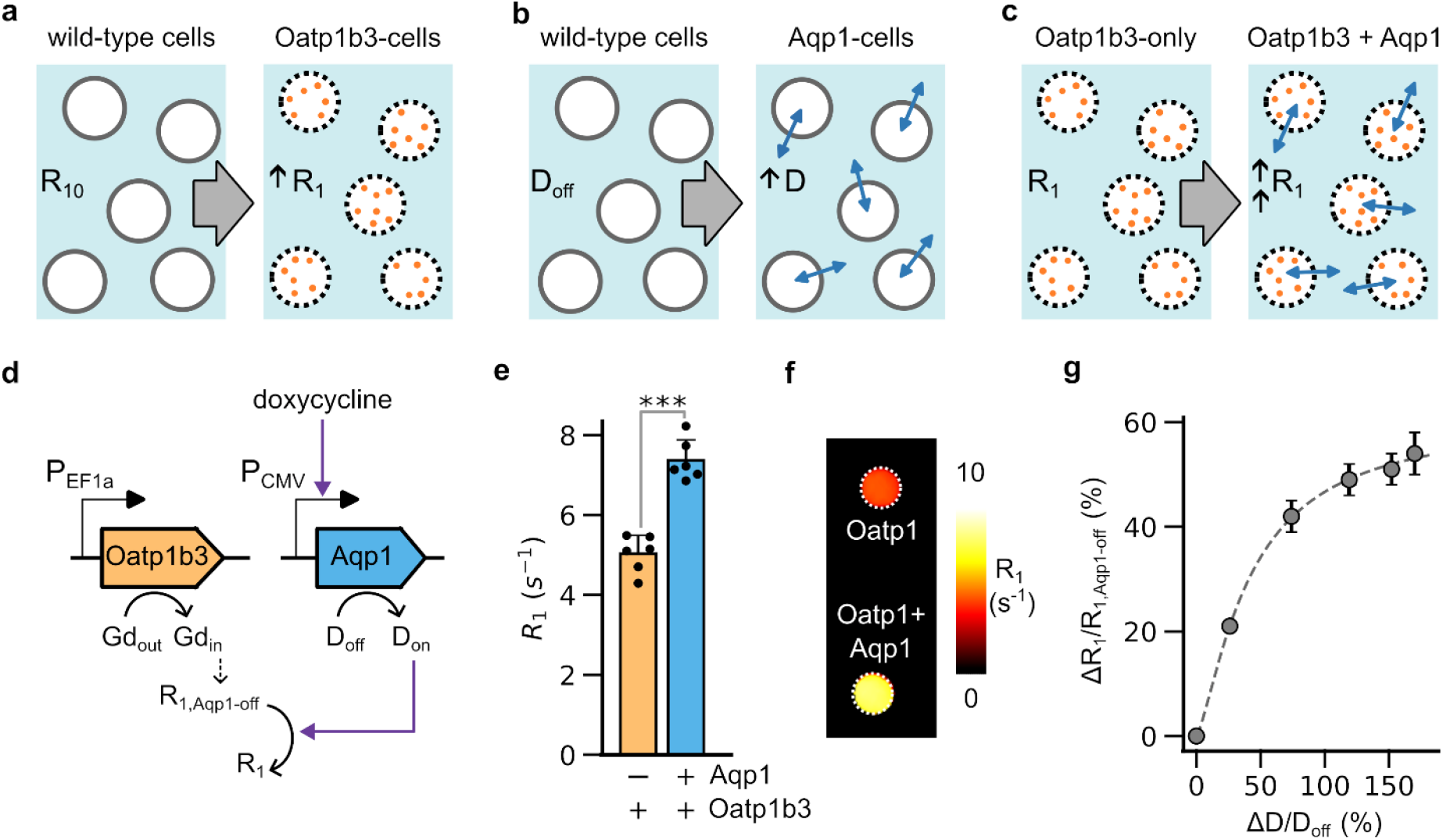
Aqp1-driven enhancement of T_1_ relaxation. **(a)** Illustration depicting the increase in the baseline T_1_ relaxation rate (R_10_) of wild-type cells (white circles) due to the expression of Oatp1b3, which promotes the intracellular uptake of Gd-EOB-DTPA (depicted as light brown, filled circles). **(b)** Illustration depicting the increase in the baseline diffusivity of cells (D_0_) due to expression of Aqp1, which facilitates transmembrane water exchange (depicted by bidirectional blue arrows). **(c)** Illustration depicting the increase in R_1_ of Oatp1b3-expressing cells due to co-expression of Aqp1, which permits intracellular Gd-EOB-DTPA to access a larger pool of water molecules via transmembrane exchange. **(d)** Schematic outline of the T_1_-VISA mechanism, which incorporates a constitutively expressed reporter gene (Oatp1b3) that promotes the uptake of Gd-EOB-DTPA; and an amplifier gene (Aqp1) that can be induced with doxycycline to enhance diffusivity (D_on_ > D_off_) and consequently elevate the T_1_-relaxation rate of cells relative to conditions where Aqp1 expression is not induced (R_1_ > R_1,Aqp1-off_). **(e)** T_1_ relaxation rate (R_1_ = 1/T_1_) of Oatp1b3-expressing cells in the absence and presence of Aqp1 expression modulated by withholding or adding doxycycline. **(f)** R_1_ maps of Oatp1b3-labeled cells in the absence and presence of Aqp1 expression. **(g)** Increase in the T_1_ relaxation rate (ΔR_1_/R_1,Aqp1-off_) of Oatp1b3-labeled cells as a function of change in diffusivity (ΔD/D_off_). To alter diffusivity, cells were treated with varying concentrations of doxycycline in the range of 0.01-1 µg/mL, which induces different levels of Aqp1 expression from the minimal CMV promoter. The dotted line represents a hyperbolic fit to the experimental data and is included to aid in the visual tracking of the trend in ΔR_1_/R_1,Aqp1-off_ vs. ΔD/D_off_. The diffusivity and relaxation measurements were conducted at ambient temperature at a field strength of 7 Tesla. Error bars represent standard deviation from multiple biological replicates. *** P-value < 0.001.

In order for Oatp1b3 to create contrast, the gadolinium centers that accumulate in Oatp1b3-expressing cells must be able to interact with water molecules. However, the native plasma membrane restricts water exchange, thereby limiting these paramagnetic interactions primarily to the water pool within Oatp1b3-expressing cells, and effectively excluding water in the extracellular volume and within surrounding cells that do not express Oatp1b3 (and therefore do not accumulate Gd-EOB-DTPA). This restriction of water access reduces effective paramagnetic relaxation enhancement, thereby limiting the minimum fraction of Oatp1b3-expressing cells that can be detected. We hypothesized that the expression of Aqp1 could overcome this limitation by providing Oatp1b3-expressing cells with access to a larger water pool, thereby allowing more water molecules to contribute to T_1_ relaxation compared to native conditions, where water exchange is limited by the native cell membrane (**Fig. 1c**). Because aquaporins enhance signals by permitting the unrestricted movement of water molecules across cellular boundaries, we refer to this mechanism as ‘T_1_-reporter visualization with increased sensitivity using aquaporins’ (T_1_-VISA) (**Fig. 1c**). Here, we describe the engineering and characterization of genetically encoded T_1_-VISAs and demonstrate their efficacy in enhancing T_1_ signals from mixed-cell populations containing low fractions of Oatp1b3-labeled cells that are otherwise undetectable based solely on reporter gene expression.

## Results and Discussion

### Aqp1 expression enhances the T_1_-relaxation rate of Oatp1b3-expressing cells by facilitating water exchange

To test the feasibility of our concept, we co-transduced Chinese hamster ovary (CHO) cell lines with lentiviral vectors engineered to express Aqp1 and Oatp1b3. The expression of Oatp1b3 was driven by a constitutive promoter, EF1α, while Aqp1 was controlled by a minimal CMV promoter containing TetR-binding sites, allowing for inducible expression using doxycycline (**Fig. 1d**). An internal ribosome entry site (IRES) was employed to co-express each gene with a fluorescent reporter, allowing for the enrichment of doubly transduced cells by fluorescence-activated cell sorting (**Supplementary Fig. 1**). We confirmed that the doxycycline treatment resulted in an increase in diffusivity (**Supplementary Fig. 2a**) but produced no significant change in T_1_ relaxation rate (R_1_ = 1/T_1_) (**Supplementary Fig. 2b)**. In contrast, Oatp1b3 expression resulted in the expected increase in R_1_ following the incubation of cells with the contrast agent, Gd-EOB-DTPA (**Supplementary Fig. 2c**). We then measured R_1_ in cells co-expressing both Oatp1b3 and Aqp1 (**Fig. 1d**) and observed a marked increase in R_1_ (ΔR_1_/R_1,Aqp1-off_ = 43 ± 3 %, mean ± s.d., *P* = 0.001, *n* = 6, 2-sided t-test) compared to cells labeled with Oatp1b3 alone, i.e., without Aqp1 expression (**Fig. 1e,f**). To rule out the possibility that the increase in R_1_ was driven by changes in the intracellular accumulation of Gd-EOB-DTPA, we used inductively coupled plasma atomic emission spectrometry (ICP-AES) to quantify gadolinium concentration, which showed that Oatp1b3-labeled cells accumulated comparable amounts of gadolinium in both Aqp1-on and off conditions (**Supplementary Fig. 3**).

**Figure 2:**
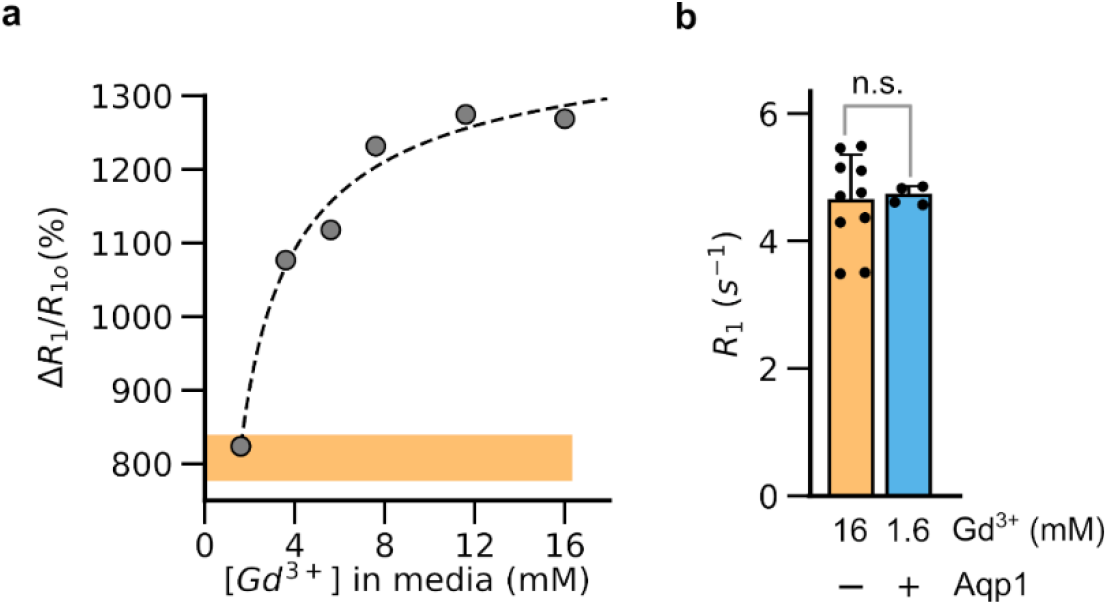
Effect of contrast agent concentration on T_1_ relaxation. **(a)** Percentage increase in R_1_ of Oatp1b3-labeled CHO cells relative to unlabeled cells (ΔR_1_/R_10_) as a function of Gd-EOB-DTPA concentration in the presence of Aqp1 expression. The dotted line represents a hyperbolic fit to the experimental data and is included to aid in the visual tracking of the trend in ΔR_1_/R_10_ vs. gadolinium. The orange band indicates the 95 % confidence interval of the increase in R_1_ (ΔR_1_/R_10_) obtained without Aqp1 expression following the incubation of cells with 16 mM Gd-EOB-DTPA. **(b)** R_1_ of Oatp1b3-expressing cells with and without Aqp1co-expression after incubation with 16 mM or 1.6 mM Gd-EOB-DTPA. Cells were incubated with the indicated Gd-EOB-DTPA concentration for 90 minutes, washed, and prepared for MRI as described in Methods. Diffusivity and relaxation measurements were conducted at ambient temperature at a field strength of 7 Tesla. Error bars represent the standard deviation from multiple biological replicates. In (a), the error bars are smaller than the data markers. n.s. P-value ≥ 0.05.

Additionally, we investigated whether co-expressing Oatp1b3 and Aqp1 as a single transcript, i.e., under the same promoter, would result in a similar Aqp1-dependent increase in R_1_. To modulate Aqp1 activity independently of Oatp1b3, we appended a destabilizing domain (DD) derived from the prolyl isomerase FKBP12 at the C-terminus of Aqp1. Our previous studies demonstrated that the Aqp1-FKBP12-DD fusion protein is constitutively destabilized, leading to a low-diffusivity off-state, but can be stabilized using a small-molecule, shield-1, which binds to the FKBP12-DD tag^42,43^. We incorporated both Oatp1b3 and Aqp1-FKBP12-DD in a single lentiviral vector separated by a ribosome-skipping 2A peptide sequence (**Supplementary Fig. 4a**), created stable CHO cell lines by viral transduction, and performed R_1_ measurements. In line with the increase in R_1_ observed with doxycycline-driven transcriptional induction of Aqp1, the modulation of Aqp1 at the post-translational level using shield-1 also results in a significant increase in R_1_ (ΔR_1_/R_1,off_ = 69.7 ± 1.3 %, P = 0.001) relative to cells that were not treated with shield-1 (**Supplementary Fig. 4b**). Collectively, these findings demonstrate the viability of the T_1_-VISAs methodology (**Fig. 1c**), which involves Aqp1 enhancing the paramagnetic relaxation of Oatp1b3-labeled cells by facilitating water exchange across the cell membrane.

Next, we examined the empirical relationship between the enhancement in water diffusivity mediated by Aqp1 (ΔD/D_off_) and the diffusion-dependent increase in relaxation rate (ΔR_1_/R_1,Aqp1-off_). To this end, we utilized cells engineered to co-express Oatp1b3 in conjunction with doxycycline-inducible Aqp1 and varied the diffusion coefficient by altering the concentration of doxycycline administered to the cells. As before, we quantified the relative increase in R_1_ compared to the baseline value obtained in the absence of Aqp1 expression. Our findings revealed that the increase in R_1_ and diffusivity exhibit an approximately hyperbolic relationship, with ΔR_1_/R_1,Aqp1-off_ tending to a plateau at high diffusion rates (**Fig. 1g**), suggesting a saturating fast-exchange regime in which the exchange rate of water molecules exceeds the Gd-EOB-DTPA induced increase in relaxation^44^.

### Aqp1 expression enables imaging using a lower concentration of contrast agent

The use of Oatp1b3 in biomedical applications requires a high dosage of Gd-EOB-DTPA to generate contrast (typically, 16 mM supplemented in growth media for in vitro imaging and >1 mmol/kg injected for in vivo applications^23,28,38–40,45^). Therefore, developing a strategy that enables Oatp1b3 to function effectively as an MRI reporter using a lower concentration of Gd-EOB-DTPA than the current experimental standard would be beneficial, particularly in situations where administering high doses of the contrast agent may be challenging or unsafe. Based on our findings above, we postulated that cells expressing both Oatp1b3 and Aqp1 could produce similar T_1_ relaxation as Oatp1b3-only cells, but using a lower dosage of Gd-EOB-DTPA. To test this hypothesis, we incubated cells co-expressing Oatp1b3 and Aqp1 with varying concentrations of Gd-EOB-DTPA ranging from 1.6-16 mM and measured the relative increase in R_1_ (ΔR_1_/R_1,Aqp1-off_) compared to Oatp1b3-only cells treated with 16 mM Gd-EOB-DTPA. As expected, ΔR_1_/R_1,Aqp1-off_ increased with increasing concentration of gadolinium (**Fig. 2a**). Notably, we found that co-expression of Oatp1b3 and Aqp1 allowed for an approximately 10-fold lower concentration of Gd-EOB-DTPA to achieve the same R_1_ as cells expressing only Oatp1b3 (**Fig. 2b**).

### Aqp1 enables detection of a small fraction of Oatp1b3-labeled cells in mixed populations

The ability to sensitively monitor small populations of genetically labeled cells is of great value in biomedical research. Therefore, we aimed to determine whether the exchange-based R_1_ amplification provided by T_1_-VISAs would enable the detection of small subsets of Oatp1b3-labeled cells in a mixed population consisting of both Oatp1b3-labeled and unlabeled cells (**Fig. 3a**). To this end, we prepared cell cultures containing varying fractions of Oatp1b3-labeled cells mixed with cells lacking Oatp1b3. Both cell types were engineered to express Aqp1 under the control of a doxycycline-inducible promoter, as previously described, thereby enabling acquisition of R_1_ measurements in the presence and absence of uniform Aqp1 expression (**Fig. 3a**). As anticipated, R_1_ increased with increasing Oatp1b3-labeled cell fraction, both in the presence and absence of Aqp1 expression (**Fig. 3b**). However, R_1_ values were consistently larger in the Aqp1-on population compared to the Aqp1-off population, as expected due to the increase in water exchange facilitated by Aqp1 expression (**Fig. 3b**). The amplification effect was lost when Aqp1 expression was switched off in Oatp1b3-expressing cells (by withholding doxycycline), further confirming that the increased access of Gd-EOB-DTPA within Oatp1b3-cells to water molecules is necessary to drive the increase in R_1_ (**Supplementary Fig. 5**). Notably, this approach achieved a nearly 4-fold increase in R_1_ even at low cell fractions (< 12 %) compared to Oatp1b3 alone i.e., without expression of Aqp1 (**Fig. 3c**). The resulting gain in sensitivity enabled the robust detection of as few as 3 % Oatp1b3-labeled cells (ΔR_1_/R_10_ = 79 ± 9 %, P = 10^-8^) that otherwise could not be detected with statistical significance based on Oatp1b3 expression alone (ΔR_1_/R_10_ = 19 ± 3 %, P = 0.06) (**Fig. 3d**).

**Figure 3:**
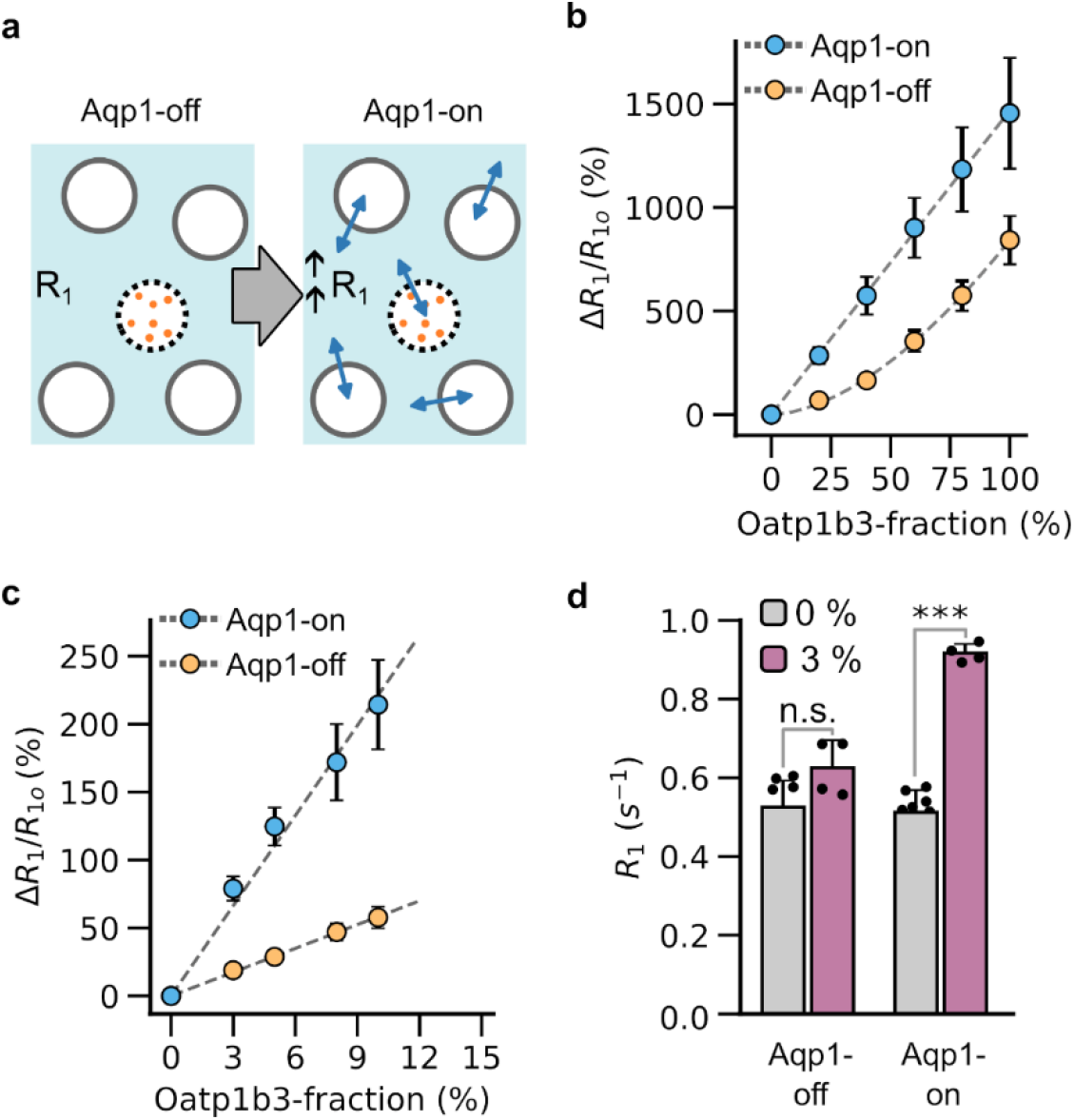
Detection of small fractions of Oatp1b3-labeled cells in mixed populations via Aqp1-driven enhancement in R_1_. **(a)** Illustration depicting a mixed-cell configuration comprising Oatp1b3-labeled cells (dotted circles with internalized Gd-EOB-DTPA labeled as filled light brown circles) and cells lacking Oatp1b3 (white circles). In the Aqp1-on state, all cells uniformly express Aqp1, enabling the amplification of R_1_ through enhanced water exchange. **(b)** Percentage increase in R_1_ (ΔR_1_/R_10_) of mixed-cell populations comprising varying fractions (20, 40, 60, 80, and 100 %) of Oatp1b3-labeled cells relative to an identically prepared cell population containing no Oatp1b3-labeled cells. Aqp1 expression was either off or induced by incubating cells with doxycycline. The dotted line represents an empirical fit to the experimental data and is included to aid in the visual tracking of the trend in ΔR_1_ /R_10_ vs. Oatp1b3-labeled cell fraction. **(c)** The percentage change in R_1_ of mixed-cell populations comprising low fractions of Oatp1b3-labeled cells (0-15 %) relative to a cell population containing no Oatp1b3-labeled cells in the presence or absence of Aqp1 expression. The dotted lines represent linear fits to the experimental data and are included to aid in the visual tracking of the functional trend in ΔR_1_/R_10_ vs. the Oatp1b3-labeled cell fraction. **(d)** R_1_ of mixed-cell populations comprising 3 % Oatp1b3-labeled cells and a population containing no Oatp1b3-labeled cells in the presence and absence of Aqp1 expression. Diffusivity and relaxation measurements were conducted at ambient temperature at 7 Tesla. Error bars represent the standard deviation from multiple biological replicates and are smaller than the data markers in some cases. *** P-value < 0.001; n.s. P-value ≥ 0.05.

## Conclusions

This study presents a novel dual gene reporter-amplifier architecture, named T_1_-VISA, for highly sensitive detection of genetically labeled cells using MRI. While previous research has demonstrated the amplification of T_1_ signals through the modulation of lipid bilayer permeability in the context of liposomes^46–48^ and MR thermometry^49,50^, the current study introduces the first fully genetic approach for exchange-based signal amplification in molecular MRI. The methodology utilizes Aqp1 and Oatp1b3, two genes that have proven effective as standalone reporters in various in vivo applications, including monitoring transcriptional activity^28,51,52^, detecting metastases^38^, tracking cell-cell communication^40^, and examining neural activity and connectivity within the mouse brain^53,54^. Compared to Oatp1b3 utilized alone, T_1_-VISA demonstrated a substantial improvement in detection sensitivity, with the ability to detect approximately 10^3^ cells per voxel. Notably, Oatp1b3 has shown success in detecting lesions estimated to comprise a few thousand cells in vivo^38^. Therefore, we anticipate that the T_1_-VISA methodology has the potential to achieve a detection limit on the order of tens of cells per voxel, which is among the highest sensitivity achieved by any genetic material for MRI to our knowledge. Furthermore, the ability to generate contrast with reduced gadolinium doses may be particularly beneficial in expanding the use of Oatp1b3 in applications where administering a large amount of contrast agent is difficult, such as for imaging in the intact brain when the reagent needs to cross the blood-brain barrier (BBB)^54^.

Initial applications of T_1_-VISAs would likely involve the localized delivery of viral vectors encoding Aqp1 expression from a constitutive promoter and Oatp1b3 from an inducible promoter to respectively allow exchange-based amplification and monitoring of transcription factor activity. However, future applications of T_1_-VISAs could greatly benefit from advances in gene delivery techniques and transgenic models, which would enable the homogeneous organ-wide expression of Aqp1 to be seamlessly integrated with the expression of Oatp1b3 from promoters that reporter on cell fate, signaling pathways, and other biological functions.

In addition to the anticipated in vivo applications, future research on T_1_-VISAs could explore the use of alternative Oatp1 variants and gadolinium-or manganese-based contrast agents^55,56^ to further enhance sensitivity and lower the contrast agent dosage. Additionally, the Aqp1-based mechanism for enhancing relaxation is not exclusive to Oatp1b3, and can be adapted to improve the performance of other classes of metal-based MRI reporters, most of which have similarly limited sensitivity. Furthermore, this approach could be used to amplify the fairly weak signal changes observed with secreted or extracellularly localized MRI reporters, such as those designed to detect neurotransmitters^57,58^ and enzyme activity^29^. Preliminary results from our lab suggest that Aqp1 expression can significantly increase the relaxation rate of cells supplemented with Gd-EOB-DTPA in the extracellular volume (**Supplementary Fig. 6**), which establishes the conceptual feasibility of using Aqp1 to amplify the relaxation enhancement induced by extracellular reporters and contrast agents. Moreover, the ability to precisely tune water diffusion using Aqp1 could offer a way to experimentally test and refine biophysical models of contrast agent relaxation, such as those based on the Bloch-McConnel-Woessner equations, which are essential for accurate interpretation of dynamic contrast-enhanced MRI in perfusion imaging. Finally, T_1_-VISAs can be harnessed to create a new class of MRI reporters for functional imaging by using aquaporin variants that regulate membrane trafficking or diffusion through the channel pore in response to biological analytes, such as calcium, pH, and enzyme activity.

In summary, we anticipate that the development of T_1_-VISAs will open up new avenues of scientific exploration in fields such as systems neuroscience, immuno-oncology, and in vivo synthetic biology where highly sensitive deep-tissue reporters are eagerly sought after for tracking biological functions in small and large animals. In practical terms, the development of T_1_-VISAs will also complement advances in gene therapies and cell-based medicine, where the ability to monitor these therapies in animals with high sensitivity will be critical to their clinical translation.

## Methods

### Molecular biology

The DNA sequences encoding Oatp1b3 (Addgene plasmid #132200) and Aqp1 were amplified using Q5® High-Fidelity 2X Master Mix (New England Biolabs, Ipswich, MA, USA) and cloned by Gibson assembly in a lentiviral transfer vector under the control of a constitutive promoter, EF1α (Addgene plasmid #60058) or a doxycycline-inducible minimal CMV promoter (Addgene plasmid #26431). EGFP and tdTomato were co-expressed with Aqp1 and Oatp1b3 respectively using an internal ribosome entry site (IRES) to facilitate the selection of stably transduced cells by fluorescence-activated cell sorting (FACS). FLAG and hemagglutinin (HA) epitope tags were added at the N-termini of Aqp1 and Oatp1b3 respectively to facilitate their detection by Western blotting, using anti-FLAG and anti-HA antibodies. The integrity all constructs was confirmed by Sanger sequencing (Genewiz, San Diego, CA, USA) or whole-plasmid nanopore sequencing (Plasmidsaurus, Eugene, OR, USA).

### Construction and packaging of lentivirus

The packaging of lentivirus was accomplished using a three-plasmid system consisting of a packaging plasmid, an envelope plasmid encoding VSV-G (to confer broad tropism), and the transfer plasmids constructed above. The three plasmids were combined in a ratio of 22 µg each for the packaging and transfer plasmids, and 4.5 µg for the envelope plasmid, and delivered to HEK293T cells by transient transfection using linear 25 kDa polyethyleneimine (Polysciences). About 24 h post-transfection, the cells were treated with sodium butyrate (10 mM) to stimulate viral gene expression. Approximately 72 h post-transfection, the supernatant was harvested and centrifuged at 500 x g for 10 min to remove cell debris. The cleared supernatant was then mixed with Lenti-X Concentrator (Takara Bio) and incubated for 24 h at 4 °C. Finally, the mixture was centrifuged at 1500 x g for 45 min at 4 °C to precipitate lentivirus. The resulting viral particles were resuspended in 0.2 mL Dulbecco’s Modified Eagle Medium (DMEM) and either immediately used for transduction of CHO cells or stored as single-use aliquots at -80 °C.

### Mammalian cell engineering

CHO cells were routinely cultured in 10-cm tissue culture grade plates using high-glucose DMEM supplemented with 10 % fetal bovine serum (R&D systems, Bio-Techne), 110 µg/mL sodium pyruvate, 100 U/mL penicillin, and 100 µg/mL streptomycin (Thermo Fisher Scientific, Waltham, MA, USA), and housed in a humidified incubator at 37 °C and 5 % CO_2_. In preparation for lentiviral transduction, CHO cells were grown to 70-80 % confluency in six-well plates, aspirated to remove spent media, and treated with purified viral particles resuspended in 0.8 mL DMEM containing 8 µg/mL polybrene (Thermo Fisher Scientific). The cells were spinfected by centrifuging the six-well plates at 1050 x g for 90 min at 30 °C and the returned to the 37 °C incubator for another 48 h. Stably transduced cells were selected by FACS using a Sony MA900 sorter and stored as cryo-stocks in the vapor phase of liquid nitrogen.

### Inductively coupled plasma atomic emission spectrometry

Following incubation of cells with Gd-EOB-DTPA for 90 min, the media was aspirated and cells were washed with sterile PBS. The cells were then treated with 0.25 % trypsin-EDTA to detach from the surface, and centrifuged at 350 x g for 5 min. The cells were washed with sterile PBS an additional 5 times, and then approximately 5 x 10^6^ cells (determined by cell counting using a C-chip™ disposable hemocytometer, Fisher Scientific) were lysed using 0.5 ml RIPA lysis buffer (Santa Cruz Biotech, Dallas, TX). Subsequently, 2 mL of 70 % (v/v) concentrated nitric acid (Fisher Scientific) was added to each sample, stirred, and heated carefully on a hot plate for approximately 24 h, inside a fume hood. After this, 1 mL of 30 % hydrogen peroxide solution (Fisher Scientific) was added to each sample, stirred, and further heated on hot plate for ∼ 24 h in a fume hood until the sample solution appeared clear and colorless. Each sample was then diluted using deionized water to reach a final concentration of 10% (v/v) nitric acid. The gadolinium concentrations in the ensuing samples were determined by Thermo iCAP 6300 Inductively coupled plasma atomic emission spectrometer (ICP-AES), using serial dilutions of a gadolinium ICP standard (Inorganic Ventures, Avantor™) for calibration.

### Magnetic resonance imaging

Prior to undergoing MRI, cells were seeded in such a manner that they reached confluence within approximately 48 h from the time of seeding. For cells harboring an inducible Aqp1 construct, doxycycline hyclate (0.01-1 µg/mL) was added to the culture media approximately 24 h after seeding. The cells were then incubated with Gd-EOB-DTPA (Eovist®, Bayer HealthCare, Germany) at concentrations ranging from 1.6-16 mM for 90 minutes before being detached from the surface by treatment with 0.25 % trypsin-EDTA (Thermo Fisher Scientific). The detached cells were then centrifuged at 350 x g for 5 minutes, washed three times with sterile phosphate buffered saline (PBS), and resuspended in 0.2 mL of sterile PBS in 0.2 plastic PCR tubes. For experiments involving mixtures of cells with and without Oatp1b3 expression, the two cell types were separately cultured and harvested as described above. The cell density was determined by cell counting using a hemocytometer, and the appropriate number of cells from each culture were mixed by gentle pipetting to achieve a desired proportion of Oatp1b3-expressing cells in the mixture. The mixed cell suspension was then centrifuged at 350 x g for 5 minutes, washed three times with sterile PBS, and transferred to 0.2 mL plastic tubes. The tubes containing the resuspended cells were centrifuged at 500 x g for 5 minutes to form a cell pellet and placed in water-filled agarose (1% w/v) molds housed in a 3D-printed MRI phantom.

All MR imaging experiments were performed in a Bruker 7T vertical-bore MRI scanner using a 66 mm diameter transceiver coil. The cell pellets were first located using a fast low angle shot (FLASH) localizer scan with the following parameters: echo time, T_E_ = 3 ms, repetition time, T_R_ = 100 ms, flip angle, α = 30 degrees, matrix size = 128 × 128, field of view (FOV) = 4.5 × 4.5 cm^2^, slice thickness = 2 mm, number of averages = 1, and total acquisition time = 12.8 s. Subsequently, a series of T_1_-weighted scans were acquired in the axial plane using a rapid acquisition with refocused echoes (RARE) pulse sequence, with the following parameters: T_E_ = 5.3 ms, flip angle, α = 90 degrees, matrix size = 128 × 128, FOV = 5.6 × 5.6 cm^2^, slice thickness = 2 mm, number of averages = 4, RARE factor = 8, total acquisition time = 13 min, and variable T_R_ = 53, 170, 302.5, 455, 635, 855, 1138, 1533, 2195, and 5000 ms. To measure cellular water diffusivity, we acquired diffusion-weighted images of cell pellets in the axial plane using a stimulated echo pulse sequence with the following parameters: T_E_ = 18 ms, T_R_ = 1000 ms, gradient duration, δ = 5 ms, gradient separation, Δ = 300 ms, matrix size = 128 × 128, FOV = 4.5 × 4.5 cm^2^, slice thickness = 2 mm, number of averages = 5, 4 effective b-values = 1115, 1794, 2294, 2794 s/mm^2^, and total acquisition time = 42 min. All images were acquired using ParaVision 6 (Bruker) and stored as DICOM files, and were analyzed using Fiji or ImageJ (National Institutes of Health). The signal intensity at a given b-value or repetition time was estimated by computing the average intensity of all voxels inside a manually drawn region of interest (ROI) encompassing the axial cross section view of a cell pellet. The relaxation time (T_1_) was calculated by fitting the growth of mean signal intensity as a function of T_R_ to an exponential function. The apparent diffusion coefficient (D) was calculated from the slope of the exponential decay in mean signal intensity as a function of effective b-value. Least-squares regression fitting was performed using the fitnlm function in Matlab (R2022b).

### Statistical analysis

Data are summarized by their mean and standard deviation obtained from *n* ≥ 3 independent biological replicates. Hypothesis testing is performed using the Student’s t-test. All tests are 2-sided and *P* value < 0.05 taken to indicate statistical significance. Quality of model-fitting (to estimate diffusivity) was ascertained based on the regression coefficient.

## Author Contributions

### Yimeng Huang

Methodology, Investigation, Validation, Formal analysis, Writing - Review & Editing. **Xinyue Chen, Ziyue Zhu**: Investigation, Validation. **Arnab Mukherjee**: Conceptualization, Methodology, Formal analysis, Resources, Data Curation, Writing - Original Draft, Writing - Review & Editing, Visualization, Supervision, Project administration, Funding acquisition.

## Acknowledgements

We thank members of the Mukherjee lab for helpful discussions. Dr. Jerry Hu (UC, Santa Barbara) is gratefully acknowledged for assistance with setting up the MRI workflow. This research was supported by the National Institutes of Health (R35-GM133530, R21-EB033989, and R01-NS128278 to A.M.). This project has been made possible in part by a grant from the Chan Zuckerberg Initiative DAF, an advised fund of Silicon Valley Community Foundation. All MRI experiments were performed at the Materials Research Laboratory (MRL) at UC, Santa Barbara. The MRL Shared Experimental Facilities are supported by the MRSEC Program of the NSF under Award No. DMR 1720256; a member of the NSF-funded Materials Research Facilities Network.

## Data availability

The data that support the findings of this study are available from the corresponding author upon reasonable request.

## Supporting Information

The following files are available free of charge: “Huang et. al. SI 2023”. The file contains supplementary figures S1-S6, depicting the fluorescence distribution of CHO cells co-transduced with Oatp1b3 and Aqp1, diffusion and R_1_ measurements of Aqp1 and Oatp1b3-expressing cells, estimation of intra-cellular gadolinium, R_1_ enhancement by post-translational stabilization of Aqp1-FKBP12-DD using shield-1, detection of small fractions of Oatp1b3-labeled cells through Aqp1-driven R_1_ enhancement, and Aqp1-driven R_1_ enhancement of Gd-EOB-DTPA treated cells.

## References

1. Helmchen, F. & Denk, W. Deep tissue two-photon microscopy. Nat Methods 2, 932–940 (2005).

2. Ntziachristos, V. Going deeper than microscopy: the optical imaging frontier in biology. Nat Methods 7, 603–614 (2010).

3. Yaghoubi, S. S., Campbell, D. O., Radu, C. G. & Czernin, J. Positron Emission Tomography Reporter Genes and Reporter Probes: Gene and Cell Therapy Applications. Theranostics 2, 374–391 (2012).

4. Minn, I. et al. Imaging CAR T cell therapy with PSMA-targeted positron emission tomography. Science Advances 5, eaaw5096 (2019).

5. Sellmyer, M. A. et al. Imaging CAR T Cell Trafficking with eDHFR as a PET Reporter Gene. Molecular Therapy 28, 42–51 (2020).

6. Keu, K. V. et al. Reporter Gene Imaging of Targeted T-Cell Immunotherapy in Recurrent Glioma. Sci Transl Med 9, eaag2196 (2017).

7. Farhadi, A., Sigmund, F., Westmeyer, G. G. & Shapiro, M. G. Genetically encodable materials for non-invasive biological imaging. Nat. Mater. 20, 585–592 (2021).

8. Farhadi, A., Ho, G. H., Sawyer, D. P., Bourdeau, R. W. & Shapiro, M. G. Ultrasound imaging of gene expression in mammalian cells. Science 365, 1469–1475 (2019).

9. Bourdeau, R. W. et al. Acoustic reporter genes for noninvasive imaging of microorganisms in mammalian hosts. Nature 553, 86–90 (2018).

10. Hurt, R. C. et al. Genomically mined acoustic reporter genes for real-time in vivo monitoring of tumors and tumor-homing bacteria. Nat Biotechnol 1–13 (2023) doi:10.1038/s41587-022-01581-y.

11. Genove, G., DeMarco, U., Xu, H., Goins, W. F. & Ahrens, E. T. A new transgene reporter for in vivo magnetic resonance imaging. Nat Med 11, 450–454 (2005).

12. Iordanova, B. & Ahrens, E. T. In vivo magnetic resonance imaging of ferritin-based reporter visualizes native neuroblast migration. NeuroImage 59, 1004–1012 (2012).

13. Cohen, B. et al. MRI detection of transcriptional regulation of gene expression in transgenic mice. Nat Med 13, 498–503 (2007).

14. Cohen, B., Dafni, H., Meir, G., Harmelin, A. & Neeman, M. Ferritin as an Endogenous MRI Reporter for Noninvasive Imaging of Gene Expression in C6 Glioma Tumors. Neoplasia 7, 109–117 (2005).

15. Gilad, A. A. et al. Artificial reporter gene providing MRI contrast based on proton exchange. Nat Biotechnol 25, 217–219 (2007).

16. Bartelle, B. B., Szulc, K. U., Suero-Abreu, G. A., Rodriguez, J. J. & Turnbull, D. H. Divalent metal transporter, DMT1: A novel MRI reporter protein. Magnetic Resonance in Medicine 70, 842–850 (2013).

17. Bartelle, B. B., Mana, M. D., Suero-Abreu, G. A., Rodriguez, J. J. & Turnbull, D. H. Engineering an effective Mn-binding MRI reporter protein by subcellular targeting. Magnetic Resonance in Medicine 74, 1750–1757 (2015).

18. Bartelle, B. B. et al. Novel Genetic Approach for In Vivo Vascular Imaging in Mice. Circulation Research 110, 938–947 (2012).

19. Ghosh, S. et al. Functional dissection of neural circuitry using a genetic reporter for fMRI. Nat Neurosci 25, 390–398 (2022).

20. Mukherjee, A., Wu, D., Davis, H. C. & Shapiro, M. G. Non-invasive imaging using reporter genes altering cellular water permeability. Nat Commun 7, 13891 (2016).

21. Patrick, P. S. et al. Development of Timd2 as a reporter gene for MRI. Magnetic Resonance in Medicine 75, 1697–1707 (2016).

22. Schilling, F. et al. MRI measurements of reporter-mediated increases in transmembrane water exchange enable detection of a gene reporter. Nat Biotechnol 35, 75–80 (2017).

23. Patrick, P. S. et al. Dual-modality gene reporter for in vivo imaging. Proceedings of the National Academy of Sciences 111, 415–420 (2014).

24. Brindle, K. M. Gene reporters for magnetic resonance imaging. Trends in Genetics 38, 996–998 (2022).

25. Gilad, A. A. et al. MRI Reporter Genes. Journal of Nuclear Medicine 49, 1905–1908 (2008).

26. Deans, A. E. et al. Cellular MRI contrast via coexpression of transferrin receptor and ferritin. Magnetic Resonance in Medicine 56, 51–59 (2006).

27. Szulc, D. A., Lee, X. A., Cheng, H.-Y. M. & Cheng, H.-L. M. Bright Ferritin—a Reporter Gene Platform for On-Demand, Longitudinal Cell Tracking on MRI. iScience 23, (2020).

28. Kelly, J. J. et al. Safe harbor-targeted CRISPR-Cas9 homology-independent targeted integration for multimodality reporter gene-based cell tracking. Science Advances 7, eabc3791 (2021).

29. Taghian, T. et al. Real-time MR tracking of AAV gene therapy with βgal-responsive MR probe in a murine model of GM1-gangliosidosis. Molecular Therapy - Methods & Clinical Development 23, 128–134 (2021).

30. Louie, A. Y. et al. In vivo visualization of gene expression using magnetic resonance imaging. Nat Biotechnol 18, 321–325 (2000).

31. Bar-Shir, A. et al. Human Protamine-1 as an MRI Reporter Gene Based on Chemical Exchange. ACS Chem. Biol. 9, 134–138 (2014).

32. Allouche-Arnon, H. et al. Computationally designed dual-color MRI reporters for noninvasive imaging of transgene expression. Nat Biotechnol 40, 1143–1149 (2022).

33. Minn, I. et al. Tumor-specific expression and detection of a CEST reporter gene. Magnetic Resonance in Medicine 74, 544–549 (2015).

34. Bar-Shir, A., Bulte, J. W. M. & Gilad, A. A. Molecular Engineering of Nonmetallic Biosensors for CEST MRI. ACS Chem. Biol. 10, 1160–1170 (2015).

35. Bricco, A. R. et al. A Genetic Programming Approach to Engineering MRI Reporter Genes. ACS Synth. Biol. 12, 1154–1163 (2023).

36. Pan, Q. et al. Organic Anion Transporting Polypeptide (OATP) 1B3 is a Significant Transporter for Hepatic Uptake of Conjugated Bile Acids in Humans. Cellular and Molecular Gastroenterology and Hepatology 16, 223–242 (2023).

37. Van Beers, B. E., Pastor, C. M. & Hussain, H. K. Primovist, Eovist: What to expect? Journal of Hepatology 57, 421–429 (2012).

38. Nyström, N. N. et al. A Genetically Encoded Magnetic Resonance Imaging Reporter Enables Sensitive Detection and Tracking of Spontaneous Metastases in Deep Tissues. Cancer Research 83, 673–685 (2023).

39. Nyström, N. N. et al. Longitudinal Visualization of Viable Cancer Cell Intratumoral Distribution in Mouse Models Using Oatp1a1-Enhanced Magnetic Resonance Imaging. Invest Radiol 54, 302–311 (2019).

40. Wang, T. et al. Visualizing cell–cell communication using synthetic notch activated MRI. Proceedings of the National Academy of Sciences 120, e2216901120 (2023).

41. Chowdhury, R. et al. Molecular imaging with aquaporin-based reporter genes: quantitative considerations from Monte Carlo diffusion simulations. 2023.06.09.544324 Preprint at 10.1101/2023.06.09.544324 (2023).

42. Yun, J. et al. Engineering ligand stabilized aquaporin reporters for magnetic resonance imaging. 2023.06.02.543364 Preprint at 10.1101/2023.06.02.543364 (2023).

43. Banaszynski, L. A., Chen, L., Maynard-Smith, L. A., Ooi, A. G. L. & Wandless, T. J. A Rapid, Reversible, and Tunable Method to Regulate Protein Function in Living Cells Using Synthetic Small Molecules. Cell 126, 995–1004 (2006).

44. Lee, J.-H. & Springer Jr., C. S. Effects of equilibrium exchange on diffusion-weighted NMR signals: The diffusigraphic “shutter-speed”. Magnetic Resonance in Medicine 49, 450–458 (2003).

45. Bhattacharyya, T., Mallett, C. L., Hix, J. M.-L. & Shapiro, E. M. MRI of OATP-expressing transplanted cells using clinical doses of gadolinium contrast agent. 2023.08.07.552326 Preprint at 10.1101/2023.08.07.552326 (2023).

46. Simon, J., Schwalm, M., Morstein, J., Trauner, D. & Jasanoff, A. Mapping light distribution in tissue by using MRI-detectable photosensitive liposomes. Nat. Biomed. Eng 7, 313–322 (2023).

47. Fossheim, S. L., Fahlvik, A. K., Klaveness, J. & Muller, R. N. Paramagnetic liposomes as MRI contrast agents: influence of liposomal physicochemical properties on the in vitro relaxivity. Magnetic Resonance Imaging 17, 83–89 (1999).

48. Permeability of liposomal membranes to water: Results from the magnetic field dependence of T1 of solvent protons in suspensions of vesicles with entrapped paramagnetic ions - Koenig - 1992 - Magnetic Resonance in Medicine - Wiley Online Library. https://onlinelibrary.wiley.com/doi/pdf/10.1002/mrm.1910230208.

49. Fossheim, S. L., Il’yasov, K. A., Hennig, J. & Bjørnerud, A. Thermosensitive paramagnetic liposomes for temperature control during MR imaging-guided hyperthermia: in vitro feasibility studies. Acad Radiol 7, 1107–1115 (2000).

50. Lindner, L. H., Reinl, H. M., Schlemmer, M., Stahl, R. & Peller, M. Paramagnetic thermosensitive liposomes for MR-thermometry. Int J Hyperthermia 21, 575–588 (2005).

51. Zhang, L. et al. Targeting visualization of malignant tumor based on the alteration of DWI signal generated by hTERT promoter–driven AQP1 overexpression. Eur J Nucl Med Mol Imaging 49, 2310–2322 (2022).

52. Li, M. et al. In vivo imaging of astrocytes in the whole brain with engineered AAVs and diffusion-weighted magnetic resonance imaging. Mol Psychiatry 1–8 (2022) doi:10.1038/s41380-022-01580-0.

53. Zheng, N. et al. A novel technology for in vivo detection of cell type-specific neural connection with AQP1-encoding rAAV2-retro vector and metal-free MRI. NeuroImage 258, 119402 (2022).

54. Li, S. et al. In vivo labeling and quantitative imaging of neuronal populations using MRI. NeuroImage 281, 120374 (2023).

55. Nyström, N. N. et al. Gadolinium-free Magnetic Resonance Imaging of the Liver via an Oatp1-Targeted Manganese(III) Porphyrin. J. Med. Chem. 65, 9846–9857 (2022).

56. McRae, S. W. et al. Development of a Suite of Gadolinium-Free OATP1-Targeted Paramagnetic Probes for Liver MRI. J. Med. Chem. 66, 6567–6576 (2023).

57. Hai, A., Cai, L. X., Lee, T., Lelyveld, V. S. & Jasanoff, A. Molecular fMRI of Serotonin Transport. Neuron 92, 754–765 (2016).

58. Shapiro, M. G. et al. Directed evolution of a magnetic resonance imaging contrast agent for noninvasive imaging of dopamine. Nat Biotechnol 28, 264–270 (2010).

