## Supplementary Information for "A dual-gene reporter-amplifier architecture for enhancing the sensitivity of molecular MRI by water exchange"

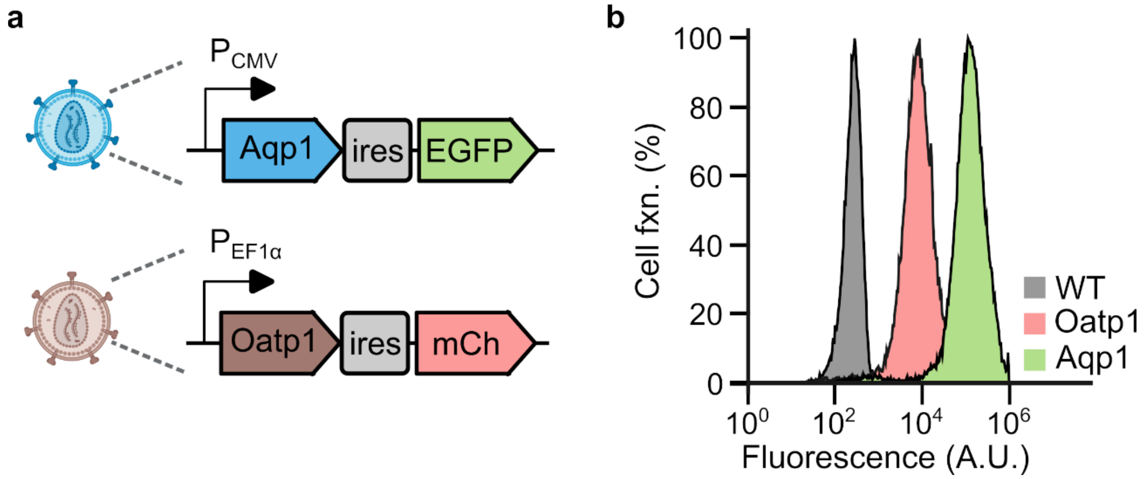


**Figure 1: Fluorescence distribution of CHO cells co-transduced with Oatp1b3 and Aqp1. (a)** Schematic of lentiviral vectors used to express reporter (Oatp1b3) and amplifier (Aqp1) genes in conjunction with fluorescent markers, enhanced GFP (EGFP) and tdTomato to enable selection of stably transduced cells by flow sorting. **(b)** Representative histogram showing fluorescence distribution of wild-type CHO cells and CHO cells co-transduced with lentiviral vectors expressing Oatp1b3-IRES-tdTomato and Aqp1-IRES-eGFP. The higher mean fluorescence of Aqp1-IRES-eGFP cells compared to Oatp1b3-IRES-tdTomato cells is attributed to the larger intrinsic brightness of eGFP relative to tdTomato.

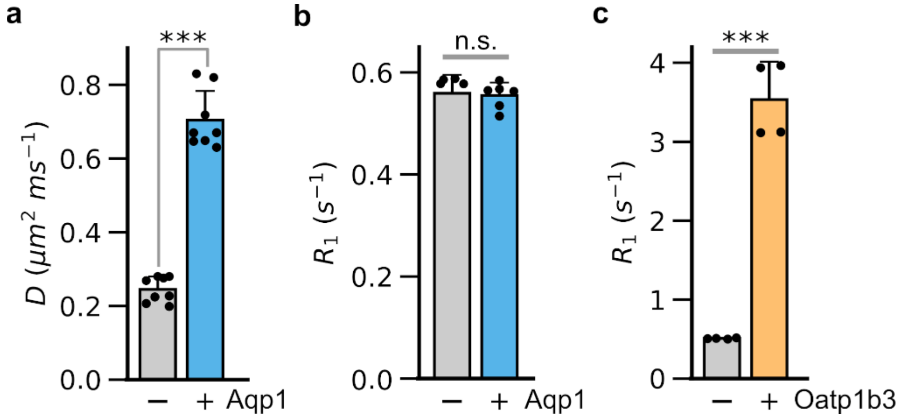


**Figure 2: Characterization of Aqp1 and Oatp1b3 functionality. (a)** Diffusivity (D) of wild-type CHO cells and CHO cells stably transduced with Aqp1. (**b**) $R_1$ of wild-type CHO cells and CHO cells stably transduced with Aqp1. **(c)** $R_1$ of wild-type CHO cells and CHO cells stably transduced with Oatp1b3. Diffusivity and $T_1$ measurements were conducted at ambient temperature using a vertical-bore 7 Tesla MRI scanner. Error bars represent the standard deviation from multiple biological replicates. ***P-value < 0.001; n.s. P-value ≥ 0.05.

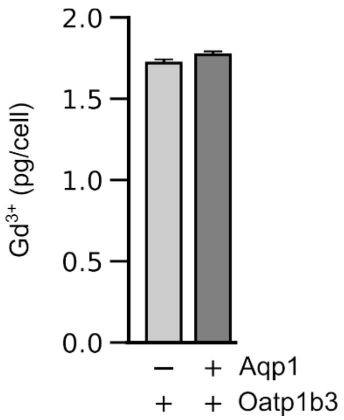


**Figure 3: Gadolinium concentration in CHO cells.** Quantification of intracellular gadolinium levels in Oatp1b3-labeled CHO cells in the absence and presence of Aqp1 (induced with 1 μg/mL doxycycline) following 90-minute incubation with Gd-EOB-DTPA. $Gd^{3+}$ levels were assessed using inductively coupled plasma atomic emission spectrometry (ICP-AES). Error bars represent the standard deviation.

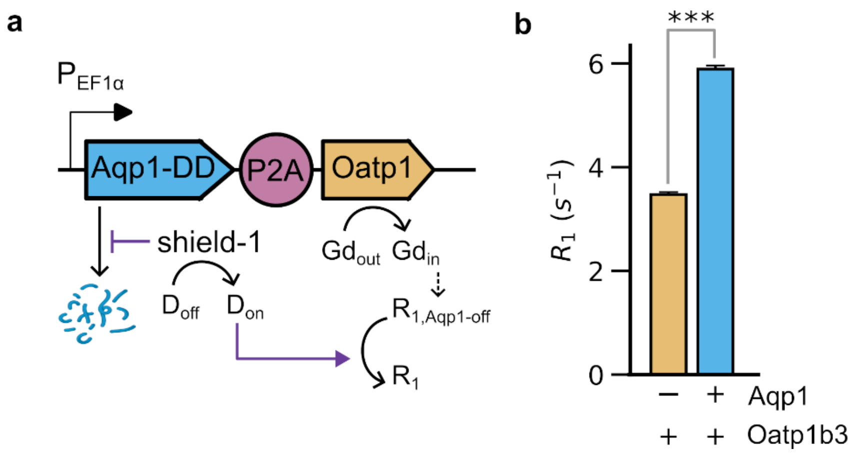


**Figure 4: Enhancement of $R_1$ by post-translational stabilization of Aqp1. (a)** Schematic representation of genetic construct engineered to co-express Oatp1b3 and Aqp1 as a single transcript. Aqp1 is tagged with a destabilizing domain (DD) derived from $FKBP12^{F36V/L106P}$, which permits post-translational modulation of Aqp1 concentration by treatment with shield-1, a small-molecule that binds to the DD and stabilizes the Aqp1-DD fusion. **(b)** $R_1$ of CHO cells stably transduced to express the single-transcript construct with and without stabilization of Aqp1 through shield-1 treatment. Error bars represent the standard deviation.

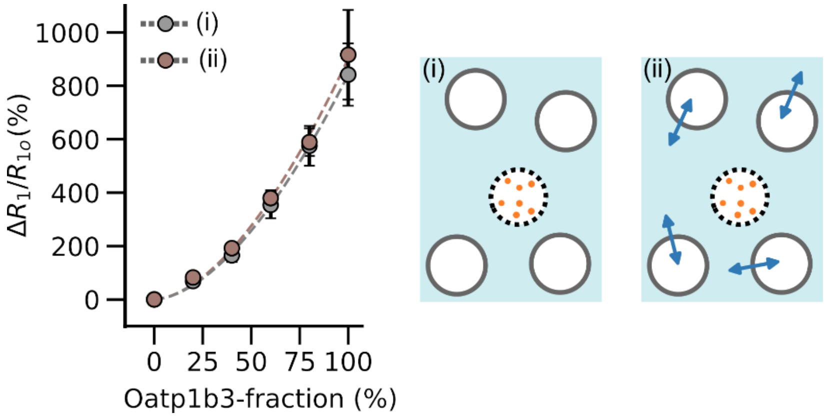


**Figure 5: Detection of small fractions of Oatp1b3-labeled cells through Aqp1-driven $R_1$ enhancement**. Percentage change in $R_1$ of mixed-cell populations corresponding to configurations (i) and (ii), relative to a homogeneous population containing no Oatp1b3-labeled cells ($\Delta R_1/R_{10}$). (i) represents mixed-cell population comprising Oatp1b3-labeled cells (dotted circle containing light brown filled circles, representing internalized Gd-EOB-DTPA) and unlabeled cells (white circles). Neither cell type expresses Aqp1. (ii) represents mixed-cell populations comprising Oatp1b3-labeled cells (that do not express Aqp1) and unlabeled cells that express Aqp1. The dotted lines represent polynomial fits to the experimental data, and are included to aid in the visual tracking of the trends in $\Delta R_1/R_{10}$ vs. Oatp1b3-labeled cell fraction. Error bars represent the standard deviation.

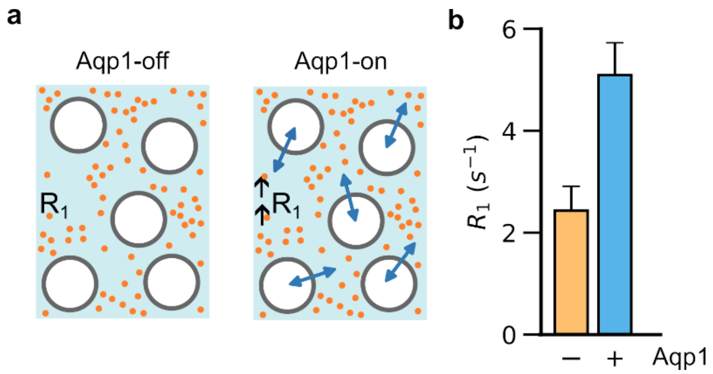


**Figure 6: Aqp1-driven $R_1$ enhancement of Gd-EOB-DTPA treated cells. (a)** Schematic outline representing a population of cells (white circles) containing Gd-EOB-DTPA (filled, light brown circles) in the extracellular space with or without water exchange (blue arrows) facilitated by Aqp1 expression. **(b)** $R_1$ of Aqp1-off and on populations of CHO cells supplemented with Gd-EOB-DTPA in the extracellular medium. Error bars represent the standard deviation.
